# Dispersal between interconnected patches can reduce the total population size

**DOI:** 10.1101/2022.04.28.489935

**Authors:** Irina Vortkamp, Christian Kost, Marita Hermann, Frank M. Hilker

## Abstract

Human activities increasingly result in a fragmentation of natural ecosystems. However, the ecological consequences of fragmentation remain poorly understood. While some studies report that fragmentation may enhance population growth, others suggest the opposite pattern. Here we investigated how habitat connectivity affects the population size of a single species when habitat patches differ in quality. We combined dispersal experiments, in which bacterial populations of *Escherichia coli* were repeatedly transferred between two qualitatively different environments, with a process-based mathematical model. Both experiments and model consistently revealed that increased dispersal between patches reduced the total population size, thus demonstrating a detrimental effect of habitat connectivity on population size. This observation could be explained with a net loss of individuals upon migration from a productive to an overcrowded patch. Our findings suggest that conservation measures, which promote movement between fragmented habitats, such as dispersal corridors or stepping stones, are potentially detrimental for some species.

## 1 Introduction

Human activities such as deforestation, agricultural land use, or urbanization have resulted in the loss and fragmentation of habitats for many species (Foley et al., 2005; Haddad et al., 2015) often with disastrous impacts on wildlife (Saunders et al., 1991; Fischer and Lindenmayer, 2007; Newbold et al., 2015). Thus, fragmentation is usually considered as being part of a degradative process that reduces biodiversity (Andren, 1994; Debinski and Holt, 2000; Margules and Pressey, 2000; Haila, 2002; Schipper et al., 2008; Pereira et al., 2010). However, a review by Fahrig (2017) reports that habitat fragmentation *per se* (i.e. the sub-division of habitats into smaller and more isolated patches without reducing the total amount of habitat) has more positive than negative effects on population occurrence, abundance, species richness, or other ecological response variables. This finding has sparked a debate about the general consequences of habitat fragmentation on the dynamics within the affected ecosystems (Fletcher et al., 2018; Fahrig et al., 2019; Miller-Rushing et al., 2019). Even for a single species, there is no clear answer to the question of whether population responses are positive or negative (Fahrig, 2017; Didham et al., 2012). Part of the problem is that the term *fragmentation* is often used as an umbrella term for processes that simultaneously reduce total habitat size, decrease habitat connectivity, and increase the length of habitat edge (Haddad et al., 2015), thus complicating the separation of these three aspects (Bunnell, 1999; McGarigal and Cushman, 2002; Hanski, 2015; Fletcher et al., 2018). While there is general agreement that habitat loss decreases biodiversity (Hanski, 2011), edge effects can be positive or negative, depending on the specific situation (Tjørve, 2010; Pfeifer et al., 2017). However, a predictive, mechanistic understanding of how habitat connectivity affects the growth of fragmented populations is lacking (Traveset and Riera, 2005; Fischer and Lindenmayer, 2007; Haddad et al., 2015). Yet, knowledge on how the connectivity between qualitatively different habitats affects the growth of a given population is crucial for choosing between conservation strategies that focus on either mitigating habitat loss or changing the spatial configuration of habitats (Villard and Metzger, 2014; Hadley and Betts, 2016).

Mathematical models can help to understand the effect of restricted movement of a population on its abundance without changing the total size of the habitat or the habitat edge. Two main outcomes have been observed: the total population density in connected habitats can be larger or smaller than the total population density of isolated patches (i.e. the sum of carrying capacities) if the habitats are qualitatively different (Freedman and Waltman, 1977; Holt, 1985; Arditi et al., 2015; Franco and Ruiz-Herrera, 2015; Wu et al., 2020). One factor that determines whether dispersal increases or decreases the population density was found to be the strength of intraspecific competition in the respective habitats as measured by the relationship between the intrinsic growth rate (*r*) and the carrying capacity (*K*) of a given population – called the *r-K relationship* (Holt, 1985; *Hendriks et al., 2005)*. *Consider two heterogeneous habitat patches, one highly productive (high K*) and one less productive (lower *K*) patch. If intraspecific competition in the more productive patch is stronger than in the less productive patch, emigration from the former to the latter can be compensated by rapid growth (high *r*) in the highly productive patch (Zhang et al., 2017). Hence, the total population density in the presence of dispersal is generally larger than in isolated patches, which is generally referred to as a *positive r-K relationship* (Fig. 1a).

**Figure 1:**
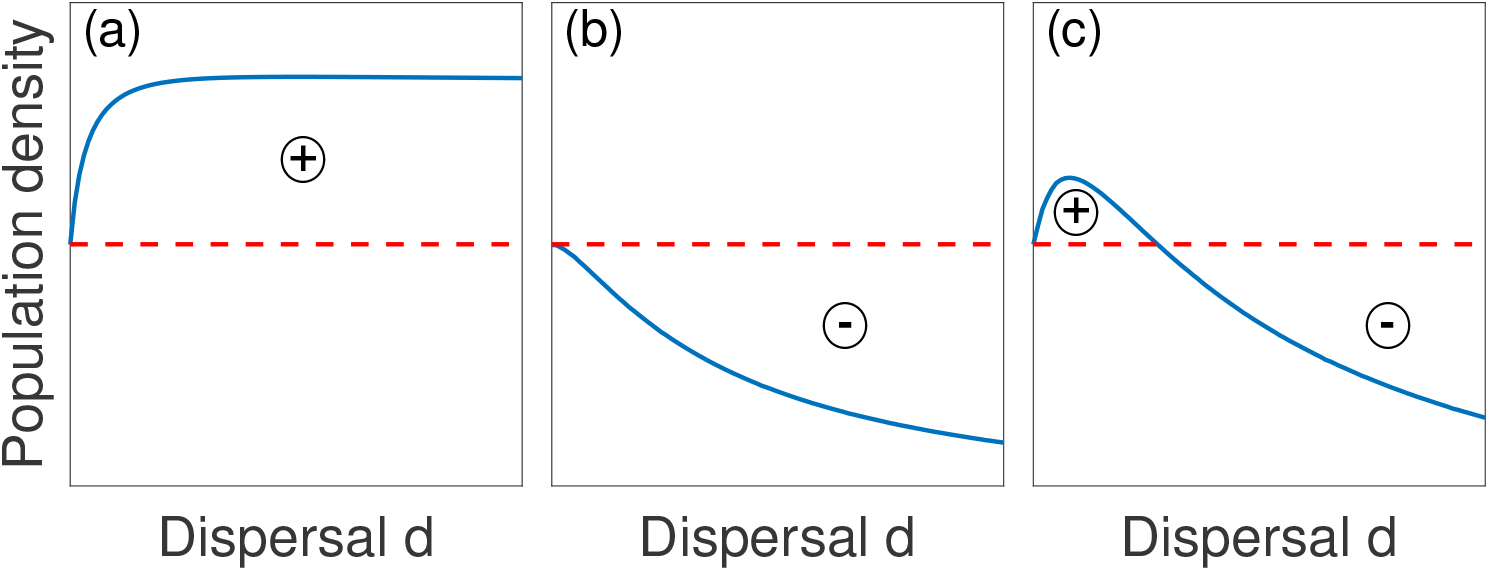
A positive r-K relationships leads to (a) positive effects of dispersal on the total population density (blue line) compared to the sum of carrying capacities (red dashed line), whereas a negative r-K relationship leads to (b) negative or (c) changing effects of dispersal on the total population density compared to the sum of carrying capacities (after Arditi et al. (2015)). Sign of effect is indicated by circled plus and minus symbols.

In contrast, if intraspecific competition is stronger in the less productive patch, losses due to emigration from the more productive patch to the patch subject to higher intraspecific competition cannot be compensated. This situation is called a *negative r-K relationship* (Zhang et al., 2017). In this case, two possible scenarios need to be distinguished: (i) increased dispersal decreases the total population density (Fig. 1b) or (ii) increased disperal non-monotonously affects the total population density (Fig. 1c) such that weak dispersal increases, while strong dispersal decreases the total population density as compared to the sum of carrying capacities.

However, it is unclear whether and in which way the parameters r and *K* are correlated in real ecological systems. A positive correlation between *r* and *K* seems plausible in habitats with fast resource renewal (Underwood, 2007; DeAngelis et al., 2020). In habitats where higher temperatures increase growth rates, higher energy needs could lead to lower carrying capacities (Underwood, 2007). When local carrying capacities are limited by available nesting sites or refuges rather than an exploitable resource, *per capita* growth rates and carrying capacities could also be uncorrelated (DeAngelis et al., 2020).

Until now it remains unclear whether highly productive habitats (high *K*) promote rapid or slow growth (high or low *r*, respectively) of a given species. Moreover, empirical evidence for positive and negative r-K relationships is even more scarce (Ives et al., 2004; Mallet, 2012; Zhang et al., 2017). Here, we combine a numerical analysis of a generic mathematical model with laboratory-based experiments with the bacterium *Escherichia coli* to investigate the long-term effect of dispersal on the population density. Our results corroborate that movement between habitats can be detrimental if the r-K relationship of two habitats is negative. Our study provides first evidence for a negative r-K relationship, which is supported by both model simulations and experimental data. These results therefore suggest that habitat fragmentation can positively affect the dynamics within fragmented ecosystems.

## 2 Material and Methods

### Model system

The long-term effect of dispersal on the population density of a single species in spatially heterogeneous habitats is investigated. We consider two habitats that differ with regard to the habitat quality, which determines growth dynamics of subpopulations (Fig. 2). A certain proportion of individuals is assumed to move between habitat patches. We systematically tested different dispersal regimes with an ascending dispersal probability to explore whether more connected or more isolated habitat patches result in higher long-term population densities.

**Figure 2:**
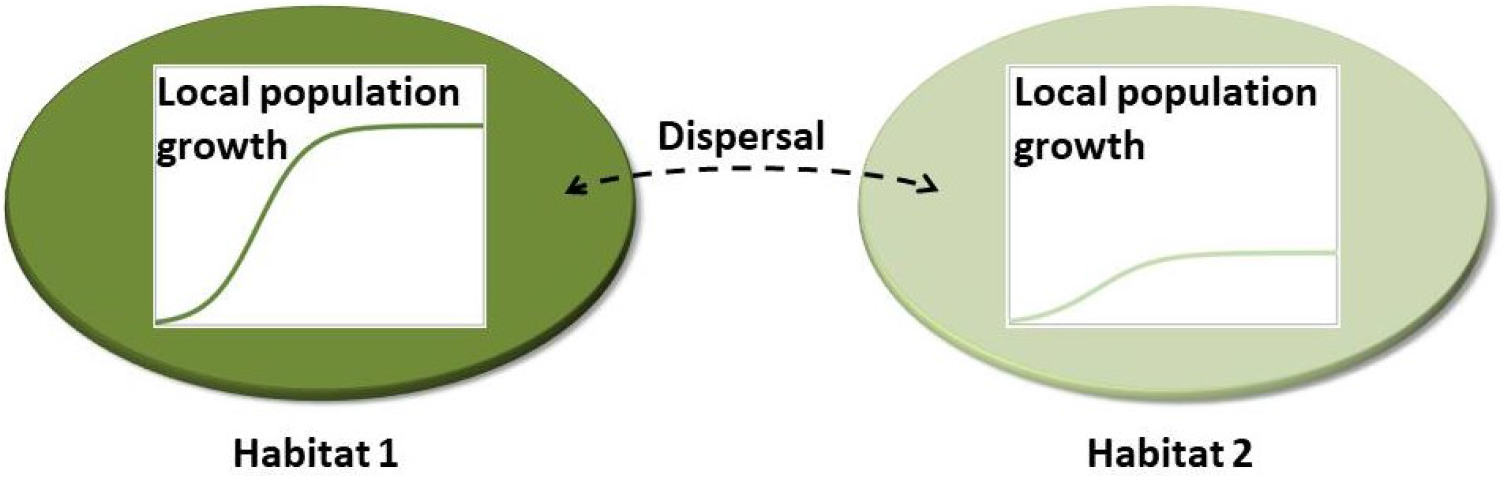
Model system used in this study. Different colors represent qualitative differences in habitat conditions leading to different local population growth dynamics over time. In the experiments, different habitats are realized by using two culture media that differ in the amount of available food resources. Dispersal is implemented by repeatedly transferring bacterial populations between habitat patches.

### Mathematical model

The growth process in the mathematical model is implemented by the Baranyi model (Baranyi and Roberts, 1994), a well-known model for bacterial population growth. This model captures both the initial lag phase and the following logistic growth. Logistic growth is typically used to model the density-dependent growth of bacterial populations (Gibson et al., 1987; Zwietering et al., 1990). A dispersal term in the model accounts for an exchange of a proportion of the population between the two habitat patches:

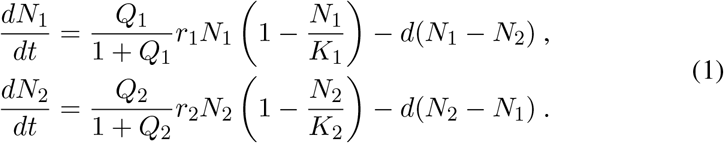

The population density in patch *i* is described by *N*_*i*_. Parameters *r*_*i*_ and *K*_*i*_ denote the intrinsic growth rate and the carrying capacity in patch *i*, respectively. In the following, patch *N*_1_ denotes the more productive habitat and *N*_2_ the less productive habitat, i.e. *K*_1_ *> K*_2_. *d* is the (symmetric) dispersal rate between patches. State variable *Q*_*i*_ accounts for the physiological state of the population, and the term 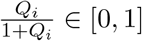 describes how well the population adapts to local habitat conditions. The term has a large impact on the transient dynamic behavior when *Q*_*i*_ is small, but no effect in the long term when *Q*_*i*_ becomes larger. The duration of the lag phase is assumed to be inversely proportional to *r*_*i*_ and related to:

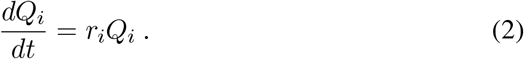

Note that *Q*_*i*_ only depends on the nutrient supply in patch *i*. It is known that the lag phase depends on many factors such as the environmental conditions, the initial cell density, the physiological stage of cells, and other factors (Swinnen et al., 2004). Following the principle of parsimony, the model neglects these aspects.

In a preliminary experiment, growth kinetics were fitted to obtain parameters *r*_*i*_ and *K*_*i*_ for the two growth environments. To this end, the log-transformed analytical solution (Baranyi and Roberts, 1994) of equations (1)-(2) without dispersal was used (see Supplementary Material A for log-transformation):

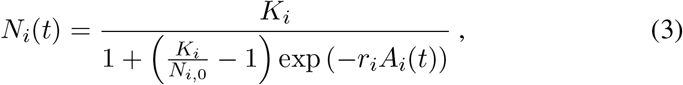

where

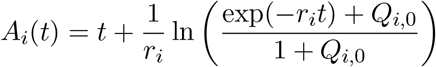

and *N*_*i*_(0) = *N*_*i*,0_ and *Q*_*i*_(0) = *Q*_*i*,0_ are initial conditions. For simplicity, in the experiments, the dispersal steps occur at discrete time steps. This discrepancy between model and experiments can lead to slightly different dynamics. However, according to Zhang et al. (2017), qualitative errors should remain small. The model was fitted to the growth kinetics experiment in order to determine the nature (positive or negative) of the r-K relationship.

### The r-K relationship

The logistic growth model makes several assumptions for population growth. First, the *per capita* growth rate, which is given by

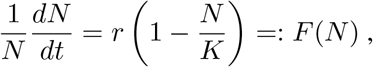

has its maximum for very low population densities at the intrinsic growth rate *r*. Second, there exists an upper limit *K* of population density due to limited resources, which is approached in the long term. Third, intraspecific competition between individuals leads to negative density-dependence. Following Holt (1985), it can be quantified by

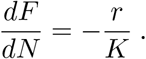

Thus, the larger *r/K*, the stronger the intraspecific competition in a habitat is (Hendriks et al., 2005). A comparison of the terms for intraspecific competition of a population in two habitats (r-K relationships) determines whether dispersal increases or decreases the overall population density at steady state (Freedman and Waltman, 1977; Holt, 1985; Arditi et al., 2015). Given a positive r-K relationship (in the following *rK*^+^), increased dispersal generally increases the total population density as compared to the sum of carrying capacities (Fig. 1a, Table 1 for parameter combinations). In contrast, for a negative r-K relationship increased dispersal always reduces the total population density (in the following *rK*^*−*^) compared to the sum of carrying capacities (Fig. 1b, Table 1 for parameter combinations) or has a non-monotonous effect (in the following *rK*^*±*^). In the latter case, weak dispersal (0 *< d < d*_crit_) leads to larger, whereas strong dispersal (*d > d*_crit_) leads to smaller total population densities compared to the sum of carrying capacities (Fig. 1c, Table 1 for parameter combinations). The critical value is *d*_crit_ = (*r*_*i*_ *− r*_*j*_)*/* [(*K*_*i*_*/r*_*i*_ *− K*_*j*_*/r*_*j*_)·(*r*_*i*_*/K*_*i*_ + *r*_*j*_*/K*_*j*_)] according to Arditi et al. (2015).

**Table 1:**
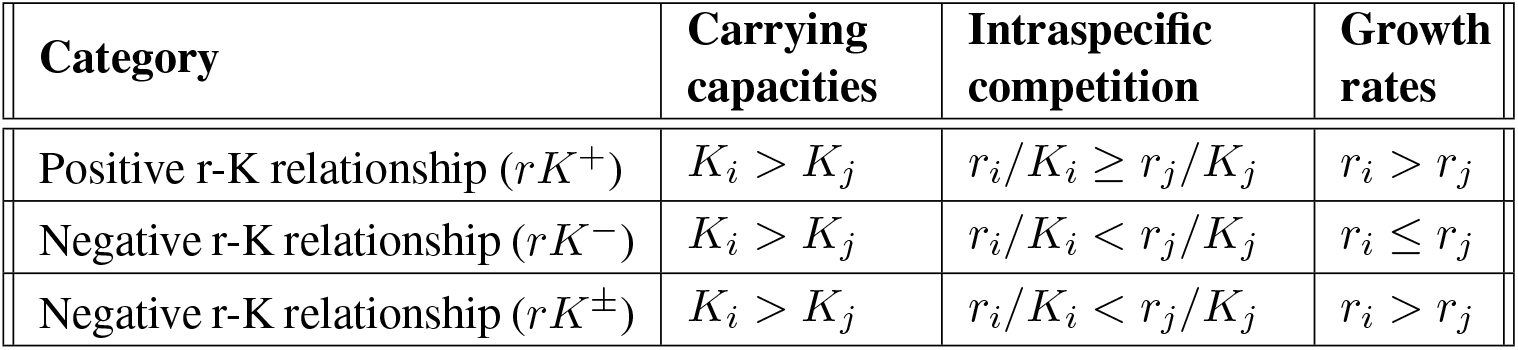
Conditions for a positive or a negative r-K relationship between two habitat patches (*i≠ j*) after Arditi et al. (2015).

| Category | Carrying capacities | Intraspecific competition | Growth rates |
| --- | --- | --- | --- |
| Positive r-K relationship ($rK^+$) | $K_i > K_j$ | $r_i/K_i \geq r_j/K_j$ | $r_i > r_j$ |
| Negative r-K relationship ($rK^-$) | $K_i > K_j$ | $r_i/K_i < r_j/K_j$ | $r_i \leq r_j$ |
| Negative r-K relationship ($rK^\pm$) | $K_i > K_j$ | $r_i/K_i < r_j/K_j$ | $r_i > r_j$ |

The r-K relationship makes qualitative predictions about the effect of dispersal on the overall population density. Hence, it can help to answer the question of whether fragmentation increases or decreases population density. However, it does not allow to draw general conclusions on how pronounced the effects are in real biological systems. This question is addressed using experimental approaches.

### Bacterial strains and cultivation conditions

For all experiments, populations of the bacterium *Escherichia coli* BW25113 (Baba et al., 2006) were used. Cells were grown in a nutrient-rich (i.e. lysogeny broth (LB) growth medium (LB Lennox, Carl Roth GmbH)) or nutrient-poor medium (i.e. LB medium diluted in 1:100 ratio with saline solution (5 gL^*−*1^ NaCl, AppliChem)). Bacterial populations were cultivated as liquid cultures at 30 °C and shaken at 200 rpm.

To initiate experiments, bacterial strains were streaked on fresh LB agar plates and incubated for 18 h or until single colonies showed sufficient size. Individual colonies were used as biological replicates to inoculate 10 mL precultures of nutrient-rich and nutrient-poor growth medium in test tubes. To minimize bacterial growth during the plating process, cultures were diluted using saline solution (5 gL^*−*1^).

### Growth kinetics

After 3 h of inoculation, precultures were adjusted to an optical density (OD) of 0.001 at 600 nm as determined via spectrophotometry in a plate reader (Spectramax M5, Applied Biosystems) and diluted 1,000-fold. 14 precultures of eight replicates – each in 1.5 mL of nutrient-rich and nutrient-poor – medium were used to start the experiment. All cultures were incubated in 96 deep-well plates (maximal volume: 2 mL, Thermo Scientific Nunc). The number of colony-forming units (CFUs) per mL culture volume was evaluated every hour (13 h in total) by drop-plating the serially-diluted culture on LB agar plates.

### Dispersal experiment

Three hours post inoculation, precultures were adjusted to an OD of 0.002 at 600 nm. For handling reasons, the initial OD was chosen to be close to the populations’ carrying capacities. Precultures were divided in four experimental groups, each consisting of one culture in 1.5 mL nutrient-rich and one culture in 1.5 mL nutrient-poor medium. Groups differed in the amount of cells that were transferred between environments: *d*_1_: no transfer between environments, *d*_2_: transfer of 300 *μ*L of the culture to the other environment, *d*_3_: transfer of 900 *μ*L of the culture to the other environment, and *d*_4_: transfer of the whole culture (1.5 mL) to the other environment (i.e. complete replacement). Four replicates of each group were used to start the experiment. Besides differences in the dispersal regime, all groups were treated in an identical way. Cultures were incubated in microtubes (max. volume: 2 mL, Eppendorf) and transferred every 1.5 h for a total of six transfers. To realize transfer of cells without mixing the media, cultures were split according to the dispersal regime (e.g. for *d*_2_: split 1.5 mL into 1.2 mL and 0.3 mL), centrifuged twice (each 2 min, 4,000 rpm), resuspended in the same volume of the corresponding medium, and reassembled. At the end of each cycle, the number of CFUs per mL culture volume was determined by drop-plating the serially-diluted culture on LB agar plates.

### Parameter estimation and statistical analysis

Parameter estimation was performed using the Matlab R2020a Curve Fitting Toolbox and lsqnonlin of the Matlab R2020a Optimization Toolbox. Cell numbers were presented on a logarithmic scale. Thus, parameters *r*_*i*_ and *K*_*i*_ were obtained by fitting the equation system in Supplementary Material A to log-transformed data generated from growth kinetics in isolated media (nutrient-rich and nutrient-poor). The initial values *Q*_*i*,0_ were obtained by fitting numerical solutions of system (1) to data from in the dispersal experiments (*d*_1_-*d*_4_).

The statistical analysis was performed using the Matlab R2020a Statistics Toolbox. Normal distribution of data was analyzed using the Lilliefors test. Homogeneity of variances was determined by applying the Brown-Forsythe test and variances were considered to be homogeneous when *P >* 0.05. Independent sample t-tests were performed against the null hypothesis that the total population density monotonically decreases with decreasing dispersal proportions. For the growth kinetics, the sample size *n* = 8 refers to the number of independent bacterial populations analyzed. In the dispersal experiment, the sample size *n* = 8 results from four independent bacterial populations, two replicates each.

## 3 Results

### Model calibration reveals negative r-K relationship

To analyze the experimental data with respect to the r-K relationships, the Baranyi model was fitted to the data generated in the growth kinetics experiments (Table 2; Supplementary Fig. B.1). Patch 1 provided better (larger *r* and *K*) growth conditions for the bacteria than patch 2. However, intraspecific competition (measured by *r/K*) appeared to be stronger in patch 2 than in patch 1. This parameter combination constitutes a negative r-K relationship with non-monotonous effect of dispersal on population density (*rK*^*±*^). Thus, weak dispersal (*d < d*_crit_) is expected to increase population density as compared to the sum of carrying capacities, whereas strong dispersal (*d > d*_crit_) is expected to decrease population density as compared to the sum of carrying capacities.

**Table 2:**
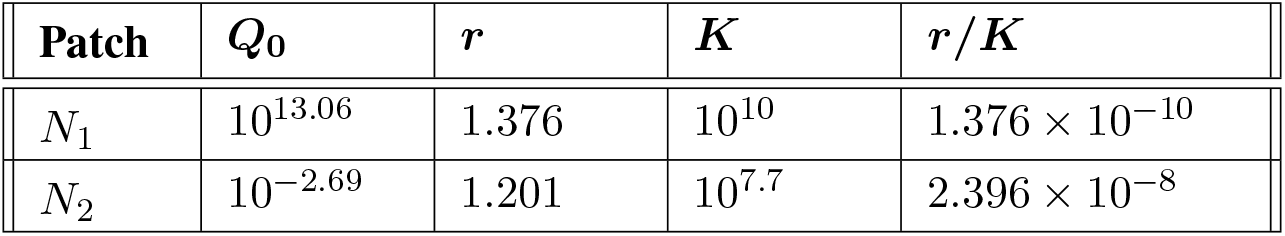
Fitted parameter values reveal a negative r-K relationship (*rK*^*±*^).

### Two populations become one

A time-series experiment, in which populations of *E*.*coli* were subjected to one of four different dispersal regimes, was performed to analyze the impact of the dispersal rate on the dynamic behavior of the system. The experimental data for the regime without transfer of cells (*d*_1_) showed a saturation over time and converged to the carrying capacity in the respective environments (Fig. 3a). The final density reached in the nutrient-rich patch (*N*_1_) was more than two orders of magnitude larger than in the nutrient-poor patch (*N*_2_). With increasing rates of dispersal between patches (*d*_2_-*d*_4_), cell densities in the two patches differed less than in the case without transfer (Fig. 3b,c,d). Small levels of dispersal between patches (*d*_2_) slightly reduced the final density in the high quality patch and increased the final density in the low quality patch (Fig. 3b). Further increasing the rate of dispersal (*d*_3_) resulted in an even stronger convergence of the population densities in the two patches (Fig. 3c). When all cells were transferred (*d*_4_), cell densities fluctuated and the amplitude of fluctuations became smaller as populations approached final population densities (Fig. 3d). For instance, the population density in *N*_2_ decreased after 1.5 h about an order of magnitude. In contrast, the decrease after 7.5 h was almost negligible, indicating that the system likely attains a stable equilibrium in the long-term.

**Figure 3:**
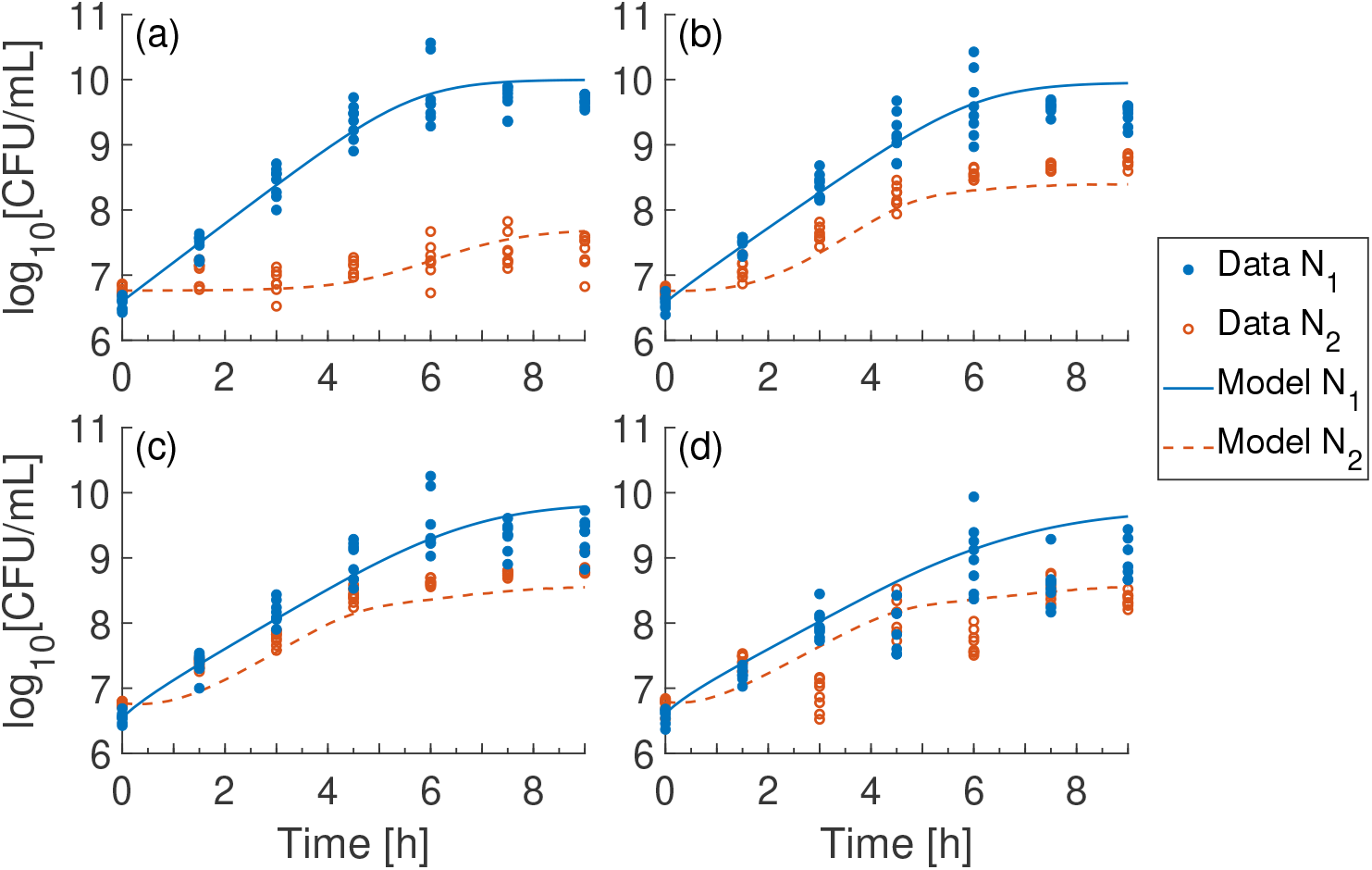
The temporal growth dynamics of the two populations were distinct when the patches were isolated (a) and became more similar with increasing dispersal (b-d). Experimental data for temporal dynamics of *E. coli* in the nutrient-rich (blue, filled) and nutrient-poor environment (orange, empty) for increasing rates of dispersal (*d*_1_-*d*_4_) (a-d respectively). Solid and dashed lines represent model predictions (see Table 2 for parameter values).

Strikingly, model solutions closely matched the experimental data (Fig. 3, blue solid and orange dashed line). The only difference between the experimentally observed data and the predictions of the model was that the latter did not show fluctuations for the highest dispersal regime (*d*_4_) (Fig. 3d). However, the final densities reached by populations of all dispersal regimes were captured well. Taken together, both model simulations and empirical data showed that population dynamics in isolated patches were rather independent from each other, yet the two patches started to converge to more similar population densities as the rate of dispersal between them increased. This dynamical behavior is known from spatial population models **??**, but has rarely been verified experimentally.

### Dispersal reduces population density in the long-run

Given that populations approached their stationary growth phase towards the end of the experiment, the final density provides information on how the different dispersal regimes affect the long-term development of total population densities (Fig. 4, boxplot). The total cell density (i.e. sum of densities of both patches) in the experiments was largest when both patches were completely isolated and continually decreased with increasing dispersal rates. Also the model confirmed the decreasing trend of the total population density for increasing dispersal rates (Fig. 4, red solid line). Since the equilibrium of population densities was not yet fully achieved at the end of the experiments, the model was additionally run until the system reached steady state, which confirmed the qualitative result (Fig. 4, red dashed line).

**Figure 4:**
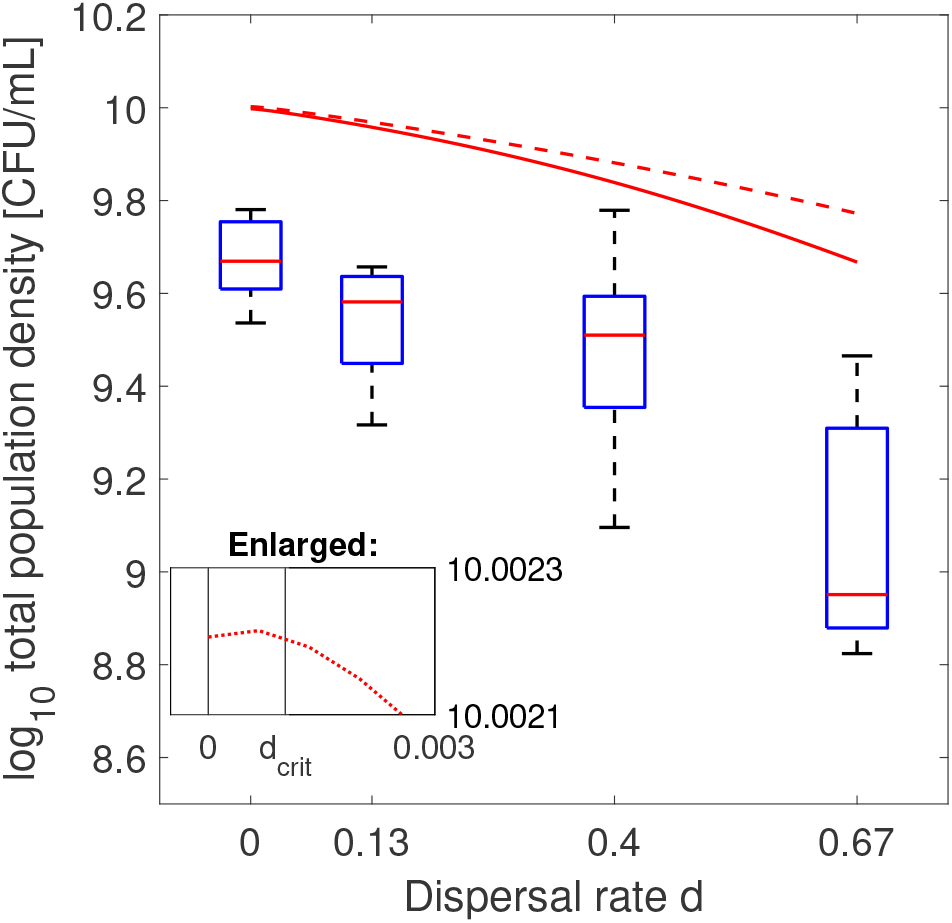
Total population density was largest in two isolated patches and decreased with increasing rates of dispersal in the experiments. The decreasing trend is significant for (*d*_1_) and (*d*_2_) (t-test: *P* = 0.0182, *n* = 8), not significant for (*d*_2_) and (*d*_3_) (*P* = 0.1859, *n* = 8), and significant for (*d*_3_) and (*d*_4_) (*P* = 0.0163, *n* = 8). The number of CFUs after 9 h of growth are shown for dispersal regimes (boxplots, *d*_1_-*d*_4_) and simulated total population densities at *t*_end_ = 9*h* (solid red line) and *t*_*∞*_ = 100*h* (dashed red line) with model (1). See Table 2 for model parameters. Experimental dispersal regimes were transformed to dispersal rates [h^*−*1^].

At first glance, the continually decreasing trend of the total population size in the experiments might not match the negative r-K relationship (case *rK*^*±*^). The fitted growth parameters in the model (Table 2) suggested that dispersal rates smaller than some critical value *d*_crit_ should increase the total population density compared to the sum of carrying capacities. Indeed, model simulations revealed a slightly larger total population density for very small dispersal rates (0 *< d < d*_crit_ *≈* 0.00102) than was observed in isolated patches (Fig. 4, inset). This result shows that even if *rK*^*±*^ predicts larger total population densities than the sum of carrying capacities for small dispersal rates, the concept does neither make any general statement about the magnitude of the critical dispersal rate *d*_crit_ nor about the magnitude of population increase. Taken together, dispersal reduced the total population density in virtually all cases analyzed.

## 4 Discussion

This study investigated the effect of dispersal between two habitats with a heterogeneous resource availability on the total population density of a single species. Combining laboratory experiments with a mathematical model revealed that dispersal reduced the overall population density compared to two isolated patches. A negative r-K relationship (*rK*^*±*^) was identified as mechanistic explanation for the observed pattern: in the nutrient-poor habitat, individuals are exposed to stronger density dependence than in the nutrient-rich habitat (Arditi et al., 2015). Consequently, as dispersal between both patches increases, a larger proportion of the overall population suffers from a stronger intraspecific competition, thus decreasing the total population size when the patches become more connected.

To our knowledge, this is the first experimental demonstration of a negative r-K relationship. Our study therefore closes the gap of previously predicted, but thus far undocumented negative r-K relationships. In combination with previous empirical studies that reported a positive correlation between population sizes and rates of dispersal (Ives et al., 2004; Zhang et al., 2015, 2017), our finding completes the empirical verification of r-K relationships as theoretically predicted by Arditi et al. (2015).

Some implicit assumptions were made in our study and will be discussed in the following. The two habitats were chosen such that the population can persist in each patch. However, some species have specific requirements regarding habitat size (e.g. home ranges), which could be undermined by fragmentation (Andren, 1994; Margules and Pressey, 2000; Hanski, 2015). Our results are valid on spatial scales where fragmentation separates populations that are connected by movement, but probably not when fragmentation occurs within individuals’ homeranges (Franklin et al., 2002; Hanski, 2011). Another assumption in our model that may need to be adapted for certain species is that population movement was modelled as symmetric passive dispersal. Animal populations, for instance, are unlikely to randomly move between heterogeneous habitats and rather prefer more suitable habitat conditions. Nevertheless, over long time scales, movement processes of both plant and animal populations can often be represented as diffusive (DeAngelis et al., 2016).

If reduced dispersal between habitats is assumed to be caused by reduced habitat connectivity, our result can be interpreted as a positive effect of fragmentation on population abundance. However, our study aims at arguing for neither positive nor negative effects of fragmentation in general (Hanski, 2011). It is well-known that the overall consequences of fragmentation are determined by a combination of factors including positive/negative edge effects, functional connectivity, and landscape complementation (e.g. Didham et al., 1996; Fischer and Lindenmayer, 2007; Villard and Metzger, 2014; Haddad et al., 2015; Fahrig, 2017). Even though habitat connectivity can both increase and decrease population densities (Weddell, 2002; Soulé et al., 2004; Fischer and Lindenmayer, 2007; Driscoll et al., 2013), the role of growth differences in heterogeneous habitats (r-K relationship) in driving fragmentation effects remained unclear so far. On that basis, our results challenge the paradigm that fragmentation is detrimental for all species (Haila, 2002; Foley et al., 2005; Fischer and Lindenmayer, 2007; Hanski, 2015). Future work should disentangle the interactive effects of habitat isolation, habitat edge, and total habitat size (Andren, 1994; Didham et al., 1996; Hobbs and Yates, 2003).

To investigate how pronounced negative r-K relationships are in other systems, we reviewed empirical studies that reported laboratory data of logistically growing populations under several types of heterogeneous environmental conditions. In most of these studies, the aim was not to investigate the effect of dispersal on the total population density, but to analyze how biotic and abiotic environmental conditions affect the growth of a given population. By analyzing the resulting data we asked whether the majority of r-K relationships reported in these studies were rather positive or negative. The results of this analysis revealed that evidence for both types of interactions is widespread (Table 3, see Supplementary Material C for further information).

**Table 3:** Empirical evidence for positive r-K relationships (*rK*^+^) and/or negative r-K relationships (*rK*^*−*^, *rK*^*±*^). x: evidence, **X**: most evidence (the frequency of this relations is three times higher than for any of the other relationships, or the number of reports for this relationship exceeds the others by over 30), otherwise no evidence. Note that studies differ with respect to the number of tested r-K relationships.

| Modeled species | Condition causing heterogeneity | $rK^+$ | $rK^-$ | $rK^\pm$ | Reference |
| --- | --- | --- | --- | --- | --- |
| *Escherichia coli* | Culture medium |  |  | x | This study |
| *Nephotettix* spp | Temperature | **X** | x | x | Valle et al. (1989) |
| *Chlamydomonas* | Mineral nutrients | x | **X** | x | Bell (1990) |
| *Anuraeopsis fissa* | Food density |  |  | x | Dumont et al. (1995) |
| Several | Toxin concentration | x | x | **X** | Hendriks et al. (2005) |
| *Chaetosiphon fragaefolii* | Host plant | x | **X** | **X** | Underwood (2007) |
| *Saccharomyces cerevisiae* | pH and dissolving oxygen | x | **X** | **X** | Salari and Salari (2017) |
| *Saccharomyces cerevisiae* | Culture medium | x |  |  | Zhang et al. (2017) |
| *Tetraselmis tetrahele* | Temperature | **X** | **X** | x | Bernhardt et al. (2018) |

In practice, it may be useful to avoid fragmentation, because fragmentation is almost always accompanied by habitat loss (Fletcher et al., 2018). However, in biological conservation much effort is invested into measures that reduce frag-mentation by bridging gaps between habitats (e.g. by dispersal corridors or stepping stones (Fischer and Lindenmayer, 2007)) or improving habitat quality of the matrix that surrounds isolated patches. Dispersal corridors allow movement between habitat fragments to increase overall habitat connectivity and, in some cases, were found to increase both species richness and population densities (Haddad and Baum, 1999; Debinski and Holt, 2000; Phillips et al., 2008). However, given that corridors can also be disadvantageous by promoting the spread of diseases or increasing predation pressure, there are good reasons to question the general benefit of corridors, at least as a default solution. Our results additionally indicate that dispersal corridors might reduce the total population size of a species if it exhibits a negative r-K relationship in the respective habitats. Conservation efforts may therefore better invest in restoration of destroyed habitat and the improvement of habitat quality than in enhancing connectivity (Villard and Metzger, 2014).

We found that species can benefit from an increased isolation between patches if overcrowding effects are stronger in the more productive than in the less productive patch. The reason for this is the net loss of individuals on the landscape level when they migrate from a productive and less crowded (‘better’) patch to a less productive and more crowded (‘worse’) patch. In this case, the population size benefits from keeping the ‘worse’ patch more separate from the ‘better’ patch. However, in this context it is important to note that both patches are habitable for the focal species, i.e. there is no source-sink relationship between both patches. Also, while reduced dispersal might be beneficial in the sense of increasing population size, it could diminish the ability of the ‘worse’ patch to recolonize the ‘better’ patch if the latter went extinct. Thus, it is also important to note that the r-K relationship is species-specific. As a consequence, under identical landscape configurations, some species can benefit from increased dispersal rates, while others might not. Taken together, our results emphasize the utility of combining mathematical modelling with laboratory-based experiments to gain fundamental mechanistic insights into the effects of habitat fragmentation (Friedman and Gore, 2017; Gokhale et al., 2018). The simple model formulation allowed rigorous mathematical proof for the effects of positive and negative r-K relationships (Arditi et al., 2015). Experimentally validating the model demonstrated a negative r-K relationship. Moreover, a literature survey revealed that the negative r-K relationship found in this study is not an exception, but occurs in many biological systems (Table 3). Hence, beneficial effects of fragmentation likely occur well beyond the model system considered in this study. Thus, our work can inform policy makers to apply appropriate conservation measures in order to mitigate potentially detrimental effects resulting from habitat fragmentation.

## Acknowledgements

CK was funded by the German Research Foundation ((SFB 944, P19: CK), (KO 3909/4-1: CK)) and Osnabrück University (EvoCell). The authors thank Jan Hendriks for the provision of the raw data to test for r-K relationships.

