## Supplementary Material for "Dispersal between interconnected patches can reduce the total population size"

#### A log-transformation

To fit model (1) from the main text without dispersal ($d = 0$) to the experimental data, it is log-transformed. Let $y_i = \log_{10}(N_i)$. Then

$$\frac{dy_i}{dt} = \frac{d\ \log_{10}(N_i)}{dt} = \frac{1}{\ln(10)} \frac{1}{N_i} \frac{dN_i}{dt} \ ,$$

where $\ln$ is the natural logarithm. Let $q_i = \log_{10}(Q_i)$ and $y_{i,\max} = \log_{10}(K_i)$. It follows:

$$\begin{aligned} \frac{dy_i}{dt} &= \frac{r_i}{\ln(10)(1 + 10^{-q_i})} \left(1 - 10^{y_i - y_{i,\max}}\right) \ , \\ \frac{dq_i}{dt} &= \frac{r_i}{\ln(10)} \ . \end{aligned}$$

The exact solution on a log-scale is

$$y_i(t) = y_{i,\max} - \log_{10} \left(1 + \frac{10^{y_{i,\max} - y_{i,0}} - 1}{\exp\left(r_i a_i(t)\right)}\right)$$

with

$$a_i(t) = t + \frac{1}{r_i} \ln \left(\frac{\exp\left(-r_i t\right) + 10^{q_{i,0}}}{1 + 10^{q_{i,0}}}\right) \ ,$$

$y_{i,0} = \log_{10}\left(N_{i,0}\right)$ and $q_{i,0} = \log_{10}\left(Q_{i,0}\right)$.

#### B Growth kinetics

The analytical solution of the log-transformed Baranyi model was fitted to the growth kinetics of *E.coli* in two isolated patches to obtain growth parameters for the respective environment (Fig. B.1). Both kinetics show the typical sigmoid curve, whereby the plot for the nutrient-rich environment (Fig. B.1, blue solid line) has not yet reached the carrying capacity visibly. However, after conducting preliminary experiments (not shown), we conclude that the curve is close to carrying capacity $K_1$ after 13 hours.

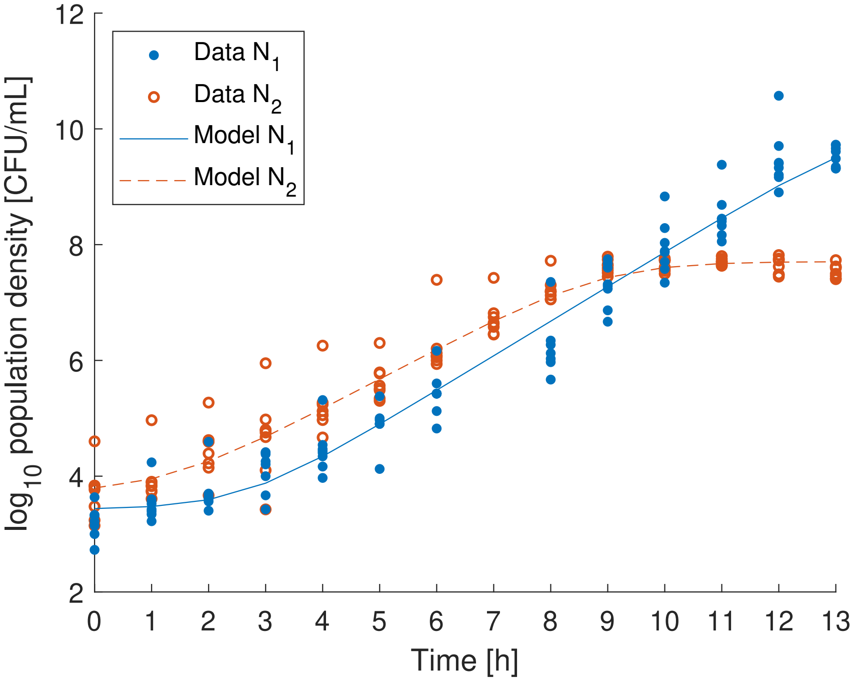


Figure B.1: Growth kinetics of *E. coli* over time (at 30°) in nutrient-rich (blue, filled dots) and nutrient-poor environment (orange, empty dots). Gap in data of $N_1$ at $t = 7$ h due to failed drop plating. Model fits were performed with the analytical solution of the log-transformed Baranyi model without dispersal. Fitted parameters: $r_1 = 1.376$, $K_1 = 10^{10}$, $r_2 = 1.201$, $K_2 = 10^{7.7}$. Fit qualities for $N_1$ and $N_2$ are $R^2 = 0.9899$ and $R^2 = 0.9937$, respectively. Sample size $n = 8$ was the same for both kinetics.

### 18 C r-K relationship

To investigate how pronounced positive and negative r-K relationships are in real biological systems, we analyzed empirical studies that report laboratory data of logistically growing populations under several types of heterogeneous environmental conditions. Where parameters were given in the references, we directly used the parameters for the analysis. Otherwise, we fitted the logistic or the Baranyi model (with lag phase) to the data to obtain parameter values. We analyzed the data by pairwise comparison of the terms for intraspecific competition ($r/K$). The outcome will be documented in upper triangular matrices in the following manner:

$$\begin{array}{c} \\ A \\ B \\ C \end{array} \begin{array}{c} \begin{array}{ccc} A & B & C \end{array} \\ \begin{pmatrix} & + & - \\ & & \pm \\ & & \end{pmatrix} \end{array}$$

In this example there are three habitats denoted by A, B, and C. Habitats A and B have a positive r-K relationship, B and C have a negative r-K relationship ($rK^{\pm}$), and A and C have a negative r-K relationship ($rK^{-}$).

#### *Nephotettix spp* (Valle et al., 1989)

Table C.1 shows mostly positive ($rK^{+}$) but also negative ($rK^{-}$ and $rK^{\pm}$) r-K relationships.

Table C.1: Fitted (logistic model) $r$ and $K$ values for different temperatures in Valle et al. (1989) to test for r-K relationship.

| **Species** | $\boldsymbol{r}$ | $\boldsymbol{K}$ | $\boldsymbol{r/K}$ | **r-K relationships** |
|---|---|---|---|---|
| *N. nigropictus* | 0.1435 | 1269 | $0.11308 \times 10^{-3}$ | $\begin{pmatrix} & + & + \\ & & \pm \\ & & \end{pmatrix}$ |
| | 0.1733 | 1315.6 | $0.13173 \times 10^{-3}$ | |
| | 0.186 | 1413.9 | $0.13155 \times 10^{-3}$ | |
| *N. virescens* | 0.1445 | 1184.8 | $0.12196 \times 10^{-3}$ | $\begin{pmatrix} & + & + \\ & & \pm \\ & & \end{pmatrix}$ |
| | 0.1726 | 1255.5 | $0.13748 \times 10^{-3}$ | |
| | 0.199 | 1580.1 | $0.12594 \times 10^{-3}$ | |
| *N. cincticeps* | 0.161 | 1428.5 | $0.11271 \times 10^{-3}$ | $\begin{pmatrix} & + & - \\ & & - \\ & & \end{pmatrix}$ |
| | 0.1809 | 1506 | $0.12012 \times 10^{-3}$ | |
| | 0.1845 | 1374.4 | $0.13424 \times 10^{-3}$ | |
| *N. malayanus* | 0.1257 | 543.3 | $0.23136 \times 10^{-3}$ | $\begin{pmatrix} & + & + \\ & & + \\ & & \end{pmatrix}$ |
| | 0.1549 | 610 | $0.25393 \times 10^{-3}$ | |
| | 0.1669 | 615.4 | $0.27121 \times 10^{-3}$ | |

#### *Chlamydomonas* (Bell, 1990)

Table C.2 shows mostly negative ($rK^{-}$) r-K relationships, few positive r-K relationships ($rK^{+}$) and one negative ($rK^{\pm}$) r-K relationship.

Table C.2: Fitted (logistic model) $r$ and $K$ values in environments with different nutrient supply in Bell (1990) to test for r-K relationship.

| $r$ | $K$ | $r/K$ | r-K relationships |
|---|---|---|---|
| 2.75 | 4.74 | 0.5794 | $-\ -\ -\ -\ -\ +\ -$ |
| 2.50 | 5.19 | 0.4819 | $-\ -\ -\ -\ -\ -$ |
| 2.79 | 4.71 | 0.5929 | $-\ -\ -\ +\ -$ |
| 2.17 | 5.74 | 0.3784 | $-\ -\ -\ -$ |
| 4.04 | 4.22 | 0.9572 | $-\ +\ +$ |
| 4.18 | 4.16 | 1.0046 | $\pm\ -$ |
| 4.89 | 5.01 | 0.9750 | $+$ |
| 4.18 | 4.36 | 0.9594 | |

#### *Anuraeopsis fissa* (Dumont et al., 1995)

Table C.3 shows negative ($rK^{\pm}$) r-K relationships.

Table C.3: Fitted $r$ (linear regression for exponential growth phase) and mean $K$ values (measured) in environments with different food supply in Dumont et al. (1995) to test for r-K relationship.

| $r$ | $K$ | $r/K$ | r-K relationships |
|---|---|---|---|
| 0.454 | 282 | $1.6 \times 10^{-3}$ | $\pm\ \pm\ \pm\ \pm$ |
| 0.4808 | 408 | $1.2 \times 10^{-3}$ | $\pm\ \pm\ \pm$ |
| 0.5344 | 666 | $0.8 \times 10^{-3}$ | $\pm\ \pm$ |
| 0.6416 | 1270 | $0.5 \times 10^{-3}$ | $\pm$ |
| 0.856 | 1989 | $0.4 \times 10^{-3}$ | |

#### Several organisms (Hendriks et al., 2005)

Meta-analysis of 95 intrinsic growth rates and carrying capacities of populations affected by toxic and other stressors. Groups of algae, rotifers, annelids, crustaceans, insects, arachnids and others were tested. Ratios of exposed and control growth parameters are compared for all species. Single parameter values are not available. Figure C.1 shows mostly negative ($rK^{\pm}$) but also positive and negative ($rK^{-}$) r-K relationships.

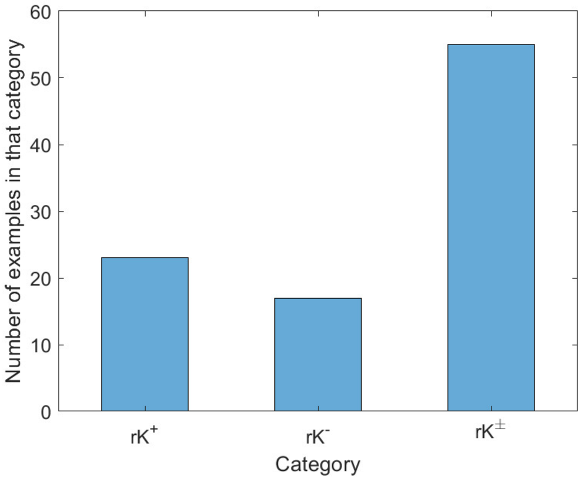


Figure C.1: $r$- and $K$-ratios in environments with and without toxin/stressor in Hendriks et al. (2005) determine the r-K relationship. Categories as defined in the main text.

#### *Chaetosiphon fragaefolii* (Underwood, 2007)

Table C.4 shows mostly negative ($rK^-$ and $rK^{\pm}$) r-K relationships, only few positive r-K relationships ($rK^+$).

Table C.4: Maximum likelihood estimates (logistic model) of $r$ and $K$ values on different host plants in Underwood (2007) to test for r-K relationship. Note that data was analyzed using webplot digitizer since no raw data was available.

| $r$ | $K$ | $r/K$ | r-K relationship |
|---|---|---|---|
| 0.176 | 8.2 | $2.146 \times 10^{-2}$ | $- \; - \; - \; - \; - \; \pm \; \pm \; - \; - \; \pm$ |
| 0.068 | 23.83 | $2.853 \times 10^{-3}$ | $+ \; + \; \pm \; \pm \; \pm \; \pm \; \pm \; \pm \; \pm$ |
| 0.099 | 25.45 | $3.890 \times 10^{-3}$ | $\pm \; \pm \; \pm \; \pm \; \pm \; \pm \; \pm \; \pm$ |
| 0.128 | 32.93 | $3.887 \times 10^{-3}$ | $- \; \pm \; \pm \; \pm \; \pm \; - \; \pm$ |
| 0.126 | 229.69 | $5.486 \times 10^{-4}$ | $\pm \; \pm \; \pm \; \pm \; - \; \pm$ |
| 0.145 | 345.79 | $4.193 \times 10^{-4}$ | $+ \; \pm \; - \; - \; \pm$ |
| 0.245 | 554.8 | $4.416 \times 10^{-4}$ | $- \; - \; - \; -$ |
| 0.198 | 575.12 | $3.443 \times 10^{-4}$ | $- \; - \; \pm$ |
| 0.141 | 723.16 | $1.950 \times 10^{-4}$ | $- \; +$ |
| 0.104 | 884.5 | $1.176 \times 10^{-4}$ | $+$ |
| 0.213 | 887.97 | $2.399 \times 10^{-4}$ | |

#### ***Saccharomyces cerevisiae*** **(Salari and Salari, 2017)**

Table C.5 shows almost exclusively negative ($rK^-$ and $rK^\pm$) r-K relationships.

Table C.5: Fitted $r$ and $K$ values (Baranyi model) in environments with different pH values and dissolved oxygen in Salari and Salari (2017) to test for r-K relationship. Note that data was analyzed using webplot digitizer since no raw data was available.

| $r$ | $K$ | $r/K$ | r-K relationship |
|---|---|---|---|
| 0.7259 | $0.3129 \times 10^{11}$ | $0.2320 \times 10^{-10}$ | $+ \; + \; \pm \; \pm \; \pm \; \pm \; - \; \pm$ |
| 1.5448 | $0.3670 \times 10^{11}$ | $0.4209 \times 10^{-10}$ | $- \; - \; - \; - \; - \; - \; -$ |
| 1.5412 | $0.3883 \times 10^{11}$ | $0.3969 \times 10^{-10}$ | $- \; - \; - \; - \; - \; -$ |
| 0.7633 | $0.4286 \times 10^{11}$ | $0.1781 \times 10^{-10}$ | $\pm \; \pm \; \pm \; - \; -$ |
| 0.8098 | $0.5048 \times 10^{11}$ | $0.1604 \times 10^{-10}$ | $\pm \; \pm \; - \; -$ |
| 1.0338 | $0.7251 \times 10^{11}$ | $0.1426 \times 10^{-10}$ | $- \; - \; -$ |
| 1.0275 | $0.7560 \times 10^{11}$ | $0.1359 \times 10^{-10}$ | $- \; -$ |
| 0.6545 | $1.1193 \times 10^{11}$ | $0.0585 \times 10^{-10}$ | $\pm$ |
| 0.7272 | $1.3256 \times 10^{11}$ | $0.0549 \times 10^{-10}$ | |

##### *Tetraselmis tetrahele* (Bernhardt et al., 2018)

Table C.6 shows mostly positive ($rK^{+}$) and negative ($rK^{-}$) r-K relationships.

Table C.6: Fitted $r$ and $K$ values (logistic model) in environments with different temperatures in Bernhardt et al. (2018) to test for r-K relationship.

| $r$ | $K$ | $r/K$ | **r-K relationship** |
|---|---|---|---|
| $0.4231$ | $0.3711 \times 10^{4}$ | $0.1140 \times 10^{-3}$ | $-$ $\pm$ $-$ $-$ |
| $0.1025$ | $1.3157 \times 10^{4}$ | $0.0078 \times 10^{-3}$ | $+$ $+$ $+$ |
| $1.4590$ | $1.7470 \times 10^{4}$ | $0.0835 \times 10^{-3}$ | $-$ $-$ |
| $0.1882$ | $1.9992 \times 10^{4}$ | $0.0094 \times 10^{-3}$ | $+$ |
| $0.2379$ | $2.0073 \times 10^{4}$ | $0.0119 \times 10^{-3}$ | |
